# Linear vector models of time perception account for saccade and stimulus novelty interactions

**DOI:** 10.1101/2020.07.13.201087

**Authors:** Amirhossein Ghaderi, Matthias Niemeier, John Douglas Crawford

## Abstract

Various models (e.g., scalar, state-dependent network, and vector models) have been proposed to explain the global aspects of time perception, but they have not been tested against specific visual phenomena like perisaccadic time compression and novel stimulus time dilation. Here, in two separate experiments (N=31), we tested how the perceived duration of a novel stimulus is influenced by 1) a simultaneous saccade, in combination with 2) a prior series of repeated stimuli in human participants. This yielded a novel behavioral interaction: pre-saccadic stimulus repetition neutralizes perisaccadic time compression. We then tested these results against simulations of the above models. Our data yielded low correlations against scalar model simulations, high but non-specific correlations for our feedforward neural network, and correlations that were both high and specific for a vector model based on identity of objective and subjective time. These results demonstrate the power of global time perception models in explaining disparate empirical phenomena and suggest that subjective time has a similar essence to time’s physical vector.

## 1 Introduction

Time perception studies generally assume that subjective or perceived time is distinct from objective time(1–3) and have explained this difference from two opposing perspectives: either from the perspective of global theories of time perception, or post-hoc explanations of specific empirical phenomena. Previous theories of time perception include linear dedicated models(4) such as internal clock models(2,3,5), and non-linear models such as state-dependent networks (SDNs)(6–8). Conversely, empirical studies have shown that various sensory and behavioral factors influence human time perception(9–13), for example, perisaccadic time compression (12–18), and dilated time for a novel visual stimulus following repeated stimuli(19–21). Perisaccadic time compression (i.e., compressed time just before and during saccades, as opposed to post-saccadic time expansion(22)) has been explained in terms of perisaccadic remapping(14,15,17) and transient cortical responses(23), whereas temporal expansion of a novel stimulus (after a series of repeated stimuli) has been attributed to a release from repetition suppression mechanisms(20,21,24). However, there is also a need to bridge these two levels of explanation (global theories vs. mechanistic models based on empirical data), because the high level models need to be empirically tested, and the low-level explanations might benefit from a unifying theory(25,26).

Here, we attempted to reconcile these approaches by 1) using empirical data (specifically, the previously unexplored interactions between perisaccadic time compression and time dilation of a novel stimulus after a series of repeated stimuli), 2) using these data to test between the predictions of the three major ‘high level’ models mentioned above (and explained below), and then 3) interpreting our findings with regard to ‘low level’ mechanistic descriptions of our findings.

In scalar timing models (Figure 1-a, Table 1), subjective time is viewed as a scalar parameter that can be generated by an internal pacemaker/accumulator structure(3) or related to levels of energy spent in the brain(13,27). Such models predict that specific time distortion effects (e.g., stimulus repetition vs. saccades) should add linearly. In contrast, SDN models(6–8,28) (Figure 1–b, Table 1) are nonlinear(8), being based on classification network training. Specifically, these models assume that time distortion effects can be coded by different connectivity patterns in specific neural networks(29). Such models can account for possible non-linear interactions between stimulus repetition and saccades. Finally, we considered a recently proposed vector model for subjective time (Figure 1-c, Table 1), based on a physical concept of time(30,31). In this model, neural time units are defined by a directional arrow and a magnitude (i.e., a vector) as in temporal physics(31–33). In this framework, simultaneous time distortions can be integrated by a neuro-information approach(30,31). For example, if the distortion effects of saccade and stimulus repetition are added to a vector of subjective time, they can then expand or compress subjective time within this vector space.

**Figure 1:**
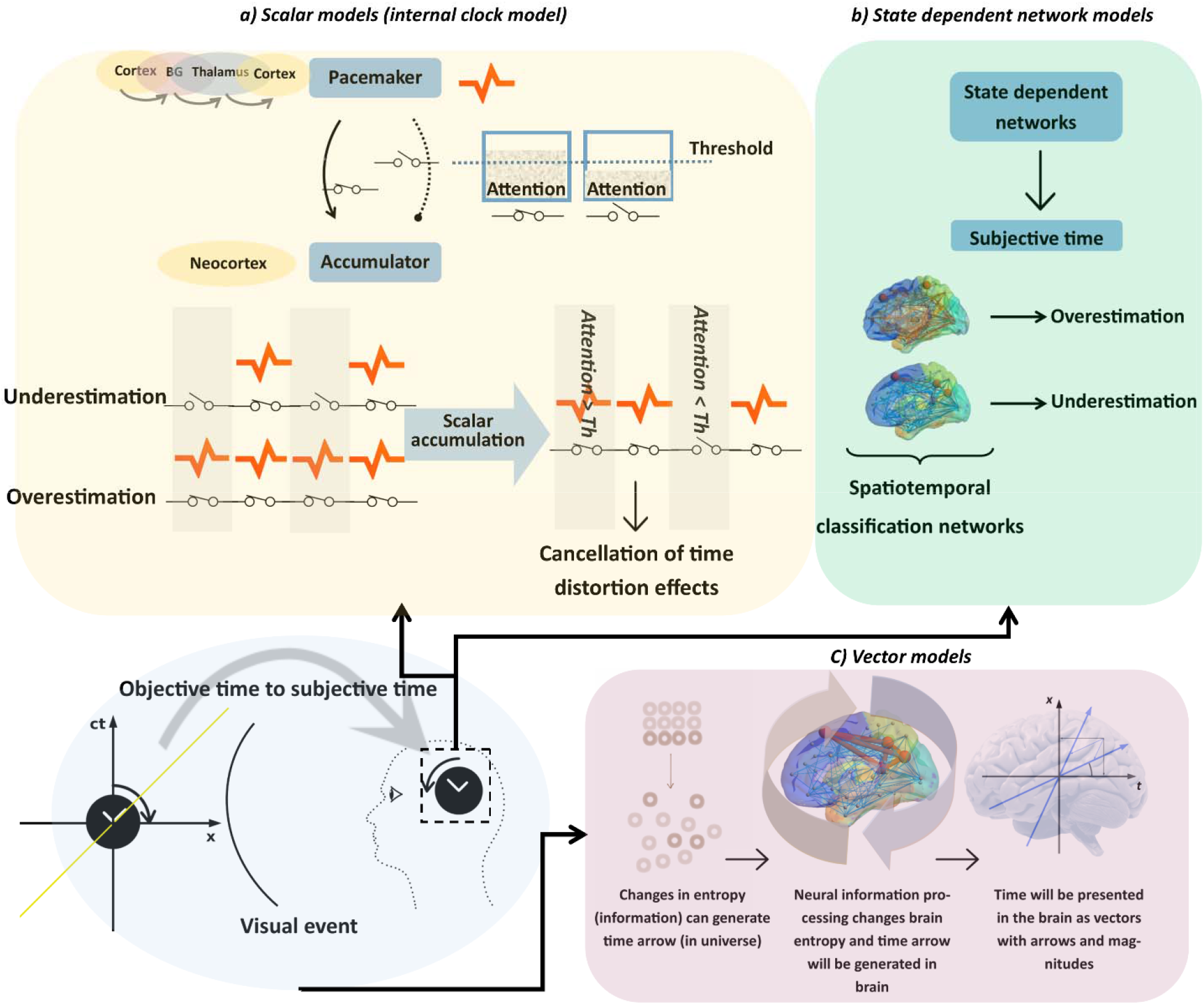
How brain can process time. The objective time that is embedded in a visual event and subjective time that is measured in the brain are characterized in two separate frameworks (universe and brain). **a) Scalar models)** *Internal clock model*: In this model, an anatomical pacemaker-accumulator structure is suggested to generate and collect scalar timing pulses. A cortico-basal ganglion-thalamocortical pathway is suggested as pacemaker(5,48,49) and different cortical regions are suggested as accumulator(5,25,48,50). In this model, attention is assumed to connect the pacemaker to accumulator as a switch(3). Usually an attention threshold is considered, and the switch will be closed when the attention is higher than this level. Otherwise, the switch is open, and accumulator cannot collect timing pulses. A higher level of attention then leads to overestimation of time whereas lower attentional allocation will lead to underestimation of time. When separate time distortion effects alter the level of attention in different directions (one increases attention and the other decreases it) the underestimation and overestimation effects cancel each other. **b) State dependent network models:** In these models different states of neural networks reflect various time perception states. The brain learns to distinguish long and short durations via Hebbian learning using a spatiotemporal classification network instead of specialized temporal structures. **c) Vector model:** A vector model explains the subjective and objective times are processed in the brain and universe via same formulization. This model describes physical parameters (e.g., entropy and speed that are involved in the measurement of objective time) are also reflected in subjective time. For example, as the entropy of the universe is increased and objective time passed, information processing in the brain increases the level of entropy in the brain and subjective time passes. In this framework, brain and universe are considered as observer and environment, respectively. Objective time can be measured in the brain based on physical time. Both timing contexts can be formulated in Minkowski space as vectors (with arrows and magnitudes).

**Table 1:** Objective and subjective time and relation between them in different time perception models.

| Model | Objective time | Subjective time | Timing rule |
| --- | --- | --- | --- |
| **Internal clock model** | Successive events | Number of neural pulses | Counting of pulses in specific brain regions can generate time. Same mechanism is suggested in Newtonian framework for physical time. |
| **Energy model** | - | Energy spent in the brain | The value of energy in the brain (a scalar parameter) can generate time. There is no identity in the physics. |
| **State dependent network model** | Successive events | Spatiotemporal classification networks | Different states of brain networks can present time. Hebbian law supports this idea. |
| **Vector model** | Physical time in Minkowski space | Physical time in Minkowski space | Physical properties of brain (i.e. entropy and speed) can measure subjective time.<br>Same mechanism is suggested in modern physics for physical time. |

To evaluate these models, we employed a repetitive stimulus / saccade / novel stimulus paradigm in human participants (Figure 2). In two separate experiments (with different stimulus durations and setups), participants judged the duration of a novel test stimulus relative to a previously viewed series of reference stimuli with a different orientation (Figure 2-a). After the reference stimuli, we cued a saccade, which commenced toward the end of the test stimulus (Figure 2-b. We then analyzed the interactions between perisaccadic time compression and repetition-induced time dilation of the test stimulus and compared these to simulations of the models described above. This yielded the novel results that time dilation after a series of repeated stimuli supersedes saccade-induced time compression, and more importantly, linear vector model simulations provided the best fits to this data (in terms of accuracy and specificity) compared to the other models of time perception. This demonstrates that global time perception models can explain specific empirical phenomena and suggests that subjective time follows the rules of objective time in a physical framework.

**Figure 2:**
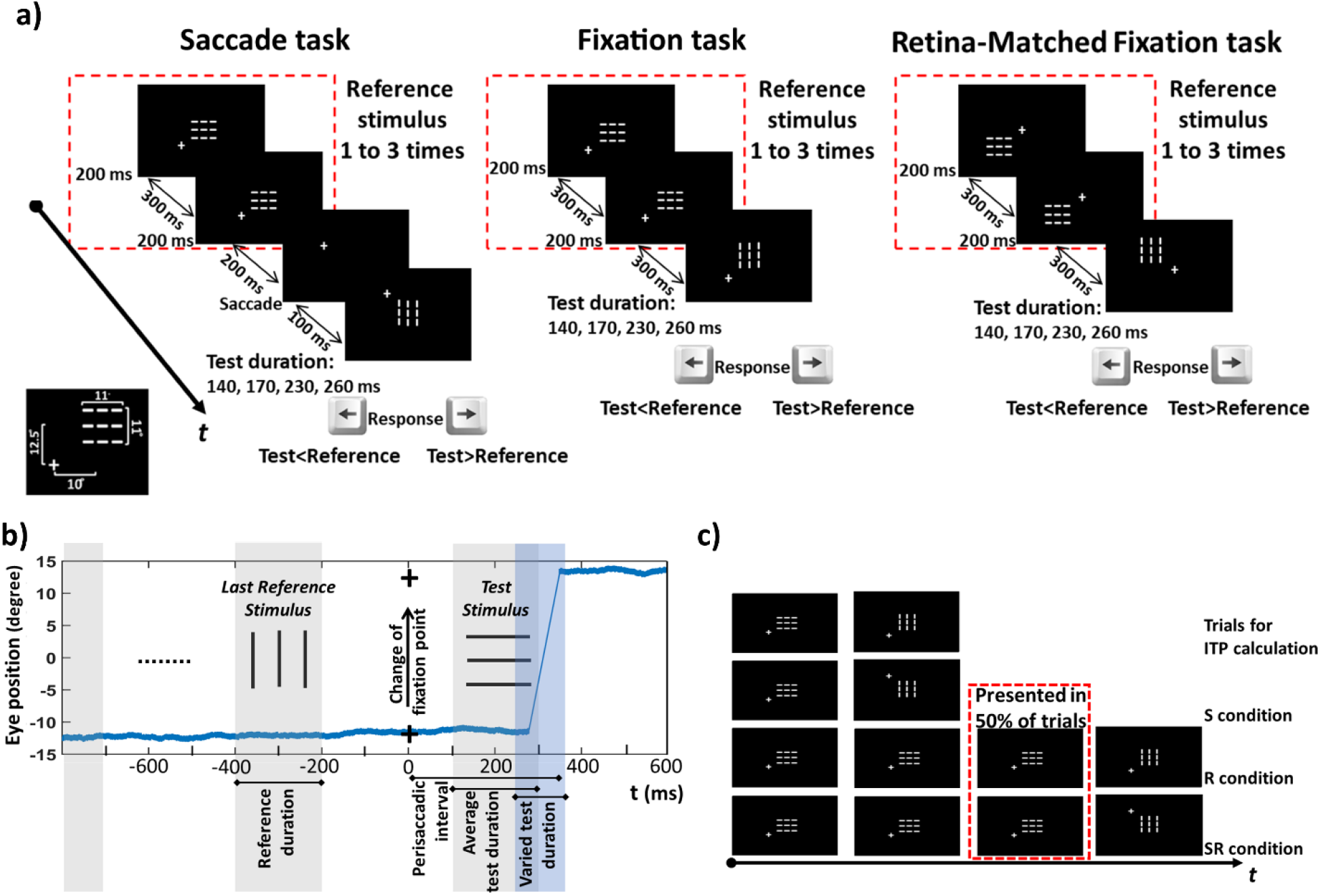
Behavioral task. **a)** Experimental design. We asked participants to judge the duration of a novel test stimulus (a horizontal or vertical grid) in comparison with the most recently presented reference stimulus (a grid with the opposite orientation. Participants fixed at one of the four corners relative to these stimuli. Three stimulus tasks (saccade, and two fixation controls) were presented in separate blocks, counterbalanced across. In each block, a series of 1-3 (randomized) 200ms reference stimuli were presented at centre, followed by the test stimulus (140, 170, 230 or 260ms, randomized) at the same or 90 deg rotated orientation. After each trial, participants judged whether the test stimulus duration was shorter or longer than the reference stimulus. Participants always began fixating 8.3 degrees diagonally from one of the four corners of the reference stimulus location. In the *Saccade Task*, the fixation point shifted up or down to the opposite corner on the same horizontal side, 100ms before test presentation, eliciting a 25 degree vertical saccade during display of the test stimulus (and causing the retinal location of the stimuli to shift). The *Fixation Task* was performed with fixed gaze and stimuli locations whereas gaze was fixed in the *Retina-Matched Fixation Task*, but the location of the test stimulus was shifted to match the retinal locations of the *Saccade Task*. **b)** Temporal evaluation and eye position in the *Saccade Task*. After presentation of the last reference stimulus (200ms), the location of fixation point was changed. Then, the test stimulus was presented (after 100ms) and eye poison was changed (after saccade latency). Presentation of the test stimulus (with variable durations) occurred during the perisaccadic interval. **c)** Separate trials were used to study the time distortion effects of saccades (S), repetition (R) and both saccades and repetition (SR). Control trials without saccades and repetition were used to calculate individual time performances (ITPs) (see details in section 2-6-1). Statistical analysis and modeling processes have been performed using this arrangement of trials.

## 2 Method

To generalize our psychophysical results, and obtain sufficient data for model testing, we include data here from 31 participants tested in two separate experiments.

### 2-1 Experiment 1

#### 2-1-1 Participants

Ten volunteers (age: mean=31.5, range between 23 and 39, 5 female) participated in the first experiment (including one of the authors; A.G.). Further data collection in this first experiment was interrupted by the pandemic. All the participants had normal/corrected to normal vision and they did not report any visual or movement disorder. All the participants signed a written informed consent. The details of experimental procedures, data collecting, and saving were described in this informed consent. This study (including the materials of informed consent) was approved by the office of research ethics (ORE) at affiliated universities. All methods in this study were performed in accordance with the Declaration of Helsinki.

#### 2-1-2 Setup

Participants sat in a dark room with their head fixed with a dental impression bar. Eye movement tracking was performed (in the *Saccade* and *Fixation Tasks*) using an Eyelink-2 system via a camera that was focused on the right pupil. Forty centimeters in front of the eyes a black display screen (1.9 × 1.4 m; luminance level of 0.015 cd/m^2^; temporal resolution: 60 Hz) presented rear-projected stimuli that were programed in C++ and build by Borland C++ version 5.02.

#### 2-1-3 Stimuli

Tree white parallel horizontal lines (thickness of line: 1°, distance between them: 4°, total size 11° ×11°) were presented as reference stimulus, while the test stimulus had the same configuration but with different orientation (vertical lines). Stimuli were presented in the right visual field (10° from the center) and a fixation point was horizontally shown at center (x=0). In the *Fixation Task*, a fixation point was randomly presented at 12.5° above or below of center (x=0°, y=±12.5°), while the location of stimuli was x=10° and y=0°. In the *Retina-Matched Fixation Task*, the fixation point was always presented at center (x=0°, y=0°) but stimuli (both of reference and test) were randomly shown 12.5° above or below of center line (x=10°, y=±12.5°). In the *Saccade Task*, the location of stimuli was same as the *Fixation Task* (x=10°, y=0°) but the location of the fixation point was randomly changed from x=0 and y=±12.5 to a symmetric location below or above an imaginary horizontal line passing through the vertical center of the screen. All the stimuli were shown on a black background.

#### 2-1-4 Trials details and experiment design

The experimental design was simplified and optimized to provide statistical power for testing the models described in subsequent Methods sections. In each task, 240 trials were presented. There were two conditions according to repetition of reference stimulus: No repetition and repetition (randomly 1 or 2 times). Duration of test stimulus was 140ms, 170ms, 230ms, or 260ms in different trials. By comparing these durations with the duration of the reference stimulus, trials were divided into longer (test duration longer than reference) and shorter trials (test duration shorter than reference). Then, in each task, 60 trials (240/(2*2)) are allocated to each condition. These conditions were presented in three different tasks: *Saccade task, Fixation task, and Retina-matched task*. In all the tasks, a reference stimulus preceded a test stimulus. Duration of the reference stimulus was always 200ms and interstimulus interval was 300ms. In the *Saccade Task*, the fixation point was moved 100ms before the presentation of the test stimulus (200ms after reference stimulus presentation). After presentation of the test stimulus participants were asked to judge which stimulus was longer, test or last reference stimulus. The procedure is illustrated in Figure 2. Trials counterbalanced location of fixation point and stimuli (i.e., in 120 trials the test stimulus was shown above and in 120 trials it was shown below the middle of the screen).

#### 2-1-5 Saccade latencies

Saccades were detected by Eyelink-2 system and recorded by a subroutine generated by visual C++ in two separate computers. The eye position signals were sent via serial port from the eye tracking computer (first device) a recording computer (second device). The task was presented on a different computer (third device) and a trigger was sent via parallel port to eye movement recording computer (second device) when the location of fixation point was changed (saccade command). The time difference between saccade command and eye movement was considered as saccade latency. The acceptable saccade latency was between 150ms and 400ms. Trials with longer or shorter latencies were removed. Since the test stimulus was always presented during saccade preparation / execution (perisaccadic interval) and vanished before fixation of the eyes at their new position (Figure 2-b), the related time distortion effect was considered to reflect perisaccadic time compression(14) rather than time expansion or chronostasis which may occur immediately following a saccade(22).

#### 2-1-6 Statistical analysis

Percentage of incorrect responses for longer and shorter trials were calculated in each condition. Incorrect responses for longer trials show underestimation of time while percentage of incorrect responses of shorter trials exhibit overestimation. We performed permutation t-tests with 5000 random shuffling for each type of these trials to evaluate the main effects of tasks and repetition. With three tasks and repetition/no-repetition of reference stimulus combinations, we had a total of 6 conditions, i.e., Saccade/No-repetition (S-NoRep), Saccade/Repetition (S-Rep), Fixation/No-repetition (Fix1-NoRep), Fixation/Repetition (Fix1-Rep), and Retina-matched Fixation/No-repetition (Fix2-NoRep), Retina-matched Fixation/Repetition (Fix2-Rep). False discovery rate (FDR) was performed to correct for multiple comparison errors. We also subtracted repetition effects from tasks to evaluate interactions between them. To perform this analysis, we compared three pairs: (*S-Rep* minus *S-NoRep*) and (*Fix1-Rep* minus *Fix1-NoRep*), (*S-Rep* minus *S-NoRep*) and (*Fix2-Rep* minus *Fix2-NoRep*), (*Fix1-Rep* minus *Fix1-NoRep*) and (*Fix2-Rep* minus *Fix2-NoRep*). Post hoc analysis, using extra permutation t-tests, were also performed among different tasks in repetition condition (i.e., S-Rep, Fix1-Rep, and Fix2-Rep). Statistical analysis was performed in MATLAB R2019a.

### 2-2: Experiment 2

#### 2-2-1 Participants

21 volunteers (age between 18 and 44, 12 female) participated in the second experiment. All participants were different from the first study. The inclusion/exclusion and ethical criteria were similar to the first experiment, except the experiment was approved by both affiliated universities.

#### 2-2-2 Setup

Stimuli were presented on a 25” LED screen (refresh rate: 144 Hz, full HD). Similar to the first experiment, participants sat in a dark room while their distance from the screen was 35cm. Fast eye movements tracking was performed (in the *Saccade* and *Fixation Tasks*) by recording electrooculography (EOG) at 2048Hz using Ag/AgCl electrodes (located at both sides of eyes) and a Biosemi amplifier. The task was programed and run using Psychtoolbox-3 (version 3.0.16) in MATLAB R2019a. Note that these data were recorded in conjunction with EEG data that are not relevant, and therefore not described or included, in the current paper.

#### 2-2-3 Stimuli

Shape and size of stimuli and fixation point were similar to the first experiment. But in the saccade task fixation point was moved horizontally from x=0, y=0 randomly to x=±12.5, y=0 (rather than vertically as in the first experiment).

#### 2-2-4 Trials details and experiment design

Participants performed a saccade and a fixation task in two different sessions (the second session was conducted one week after the first). In total 720 trials were collected in each task. In this experiment, presentation of stimuli was similar to the first experiment, except timing was changed. We set the duration of the test stimulus to 70ms and the duration of the reference to 30, 50, 90, or 110ms. The interstimulus interval was 300ms. After presentation of the test stimulus participants judged which stimulus was longer, test or last reference stimulus.

#### 2-2-5 statistical analysis

As in the first experiment, percentage of incorrect responses for longer/shorter trials were obtained in each condition to evaluate underestimation/overestimation of time. Permutation t-tests with 5000 random shuffling for each type of trials were performed and the main effects of tasks and repetition were compared. In this experiment, two tasks were presented, and we had a total of 4 conditions, i.e., Saccade/No-repetition (S-NoRep), Saccade/Repetition (S-Rep), Fixation/No-repetition (F-NoRep), and Fixation/Repetition (F-Rep). We performed false discovery rate (FDR) to correct for multiple comparison errors. Furthermore, we subtracted repetition effects from tasks to evaluate interactions between them comparing two pairs: (*S-Rep* minus *S-NoRep*) and (*F-Rep* minus *F-NoRep*). We also performed extra permutation t-tests as post hoc analysis to compare different tasks in the repetition condition (i.e., S-Rep and F-Rep). Statistical analysis was performed in MATLAB R2019a.

### 2-5 Exclusion criteria

The exclusion criteria for rejected trials were as follows: 1) Trials with saccades during standard/test stimuli presentations during fixation intervals. 2) Saccade trials with late (later than 400ms after fixation-point movement) or early (earlier than 150ms after fixation-point movement) saccade execution. 3) Trials with no responses. 4) Trials where participants were affected by any unintended external stimulus (e.g., extra noises, talking, movement etc.) during the trial presentation. Based on these criteria, the percentage of data exclusion was less than 30 % of all trials for each participant (mean=16.8%, SD=8.79%). Trial rejection was performed by self-developed subroutines in MATLAB R2019a.

### 2-6 Models

#### 2-6-1 Preparing data for models

We calculated individual timing performance (ITP) as a person’s ability to perceive time in a given condition relative to their ability in the control conditions (i.e., conditions with the minimum of distortion effects). To this end, we first obtained the percentage of trials that were correctly judged by each participant in each condition. The percentage of correctly judged trials in the conditions with no time distortion effects (i.e., repetition and saccade) was considered as pure time (*t_pure_*). These trials were obtained from both *Fixation Tasks* where the reference stimulus was presented without repetition (when there were neither repetition nor saccade effects on timing judgment) (See Figure 2-c). Equivalently, the *t_S_, t_R_, t_SR_* were defined as the percentage of trials that were correctly judged in the saccade, repetition, and saccade/repetition conditions, respectively. Next, we calculated the ratio of *t_S_, t_R_, t_SR_* in comparison to *t_pure_* and the ITP for each condition was defined as:

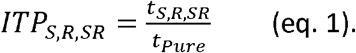

The ITPs were calculated for both types of trials (i.e., trials with shorter or longer duration of test stimulus in comparison to the duration of reference stimulus). ITP = 1 shows the duration judgment in presence of time distortion effects is similar to the duration judgment in the conditions without distortion effects. ITPs > 1 signify the time distortion effects increased accuracy of test stimulus duration judgment, whereas ITPs < 1 signify the distortion effects reduced accuracy to judge to duration of test stimulus relative to the reference stimulus. ITPs were calculated for each participant using MATLAB R2018a.

Then, we tested three different models (linear scalar model, linear vector model and state dependent network model) based on previous hypotheses about time perception (Figure 1, 3). In all three models, the same two inputs *ITP_R_, ITP_S_*, and same output *ITP_SR_* were utilized. Three criteria were selected to test the models. 1) Sensitivity (outputs of the model should correlate with the actual behavioral data). 2) Specificity (i.e., for specific model predictions these correlations should disappear when the data are randomly shuffled, whereas correlations due to overfitting the data should persist). 3) Generality (models should be compatible with different experimental conditions and various type of time judgments). These criteria were determined through linear correlation analysis and calculating root mean square error (RMSE) between output of models and original/shuffled data.

**Figure 3:**
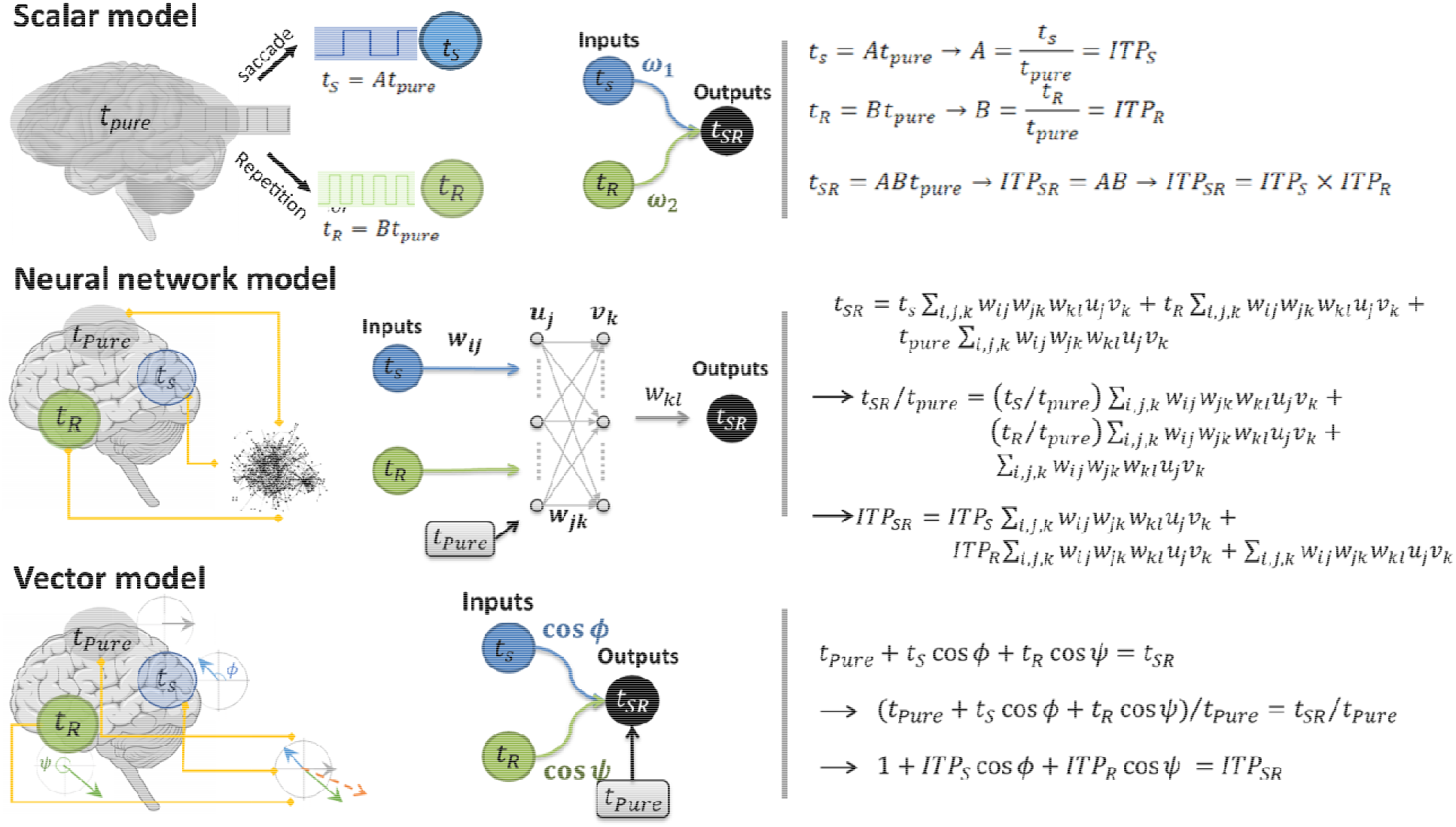
Three models have been shown schematically and mathematically. The were defined as the percentage of trials that were correctly judged in the saccade, repetition, and saccade/repetition conditions respectively and *ITP* was defined based on Eq. 1. In all the models, inputs are the *ITP* in *S* and *R* conditions and output is *ITP* in *SR* condition. **Above)** In the linear scalar timing model, an internal clock generates clock pulses and distortion effects (saccade and repetition) change the rate of pulses. as two scalar parameters with an accumulative property. Mathematically (right side), in the scalar timing model one can assume that both time distortions effects (i.e. saccade (*t_s_*) and repetition (*t_R_*)) can be added (by applying arbitrary coefficients or weights) and this scalar summation is equal to perceived time when both time distortion effects present (i.e. *t_SR_*). Since the ITP is achieved from our experimental results, both sides of equation were divided by *t_pure_* (second line). Then based on eq. 1, *t_S,R,SR_* were replaced by *ITP_S,R,SR_* (third line). All mathematical procedures in this approach are linear. **Middle)** In the nonlinear neural network model, time may be generated by state dependent networks. These networks were simulated by feedforward multilayer perceptrons. The synaptic weights are associated with a prior knowledge and based on these synaptic weights, network generates responses for new situations. Mathematically (right side), each time distortion effect (*t_s_* and *t_R_*) multiplies by several synaptic weights between layers and adds to the time in absence of distortion effects (*t_pure_*). This summation is equal to time distortion effects in the saccade and repetition condition (*t_SR_*). As in the scalar timing model, both sides were dived by *t_pure_* and according to eq. 1, 1, *t_S,R,SR_* were replaced by *ITP_S,R,SR_* **Below)** In the linear vector timing model, time units present as vectors in different angles. The angles may be defined by information processing, entropy, or speed of neural processing. This model can present a computational framework to explain how time units can be measured or integrated in the brain. Mathematically (right side), time distortion effects (*t_S_* and *t_R_*) consider as two vectors and can be added to the vector of pure time (*t_pure_*) via vector subtraction/addition operations. The vector sum is performing by applying the *cos* of the angles between the vectors and this summation is supposed to equal to perceived time in saccade and repetition condition (*t_SR_*) (the first line). Then, both sides were divided by *t_pure_* to replace *t_R,S_* with *ITP_R,S_* (the second and third rows).

#### 2-6-2 Description of models

The linear scalar model was implemented based on a wide range of studies that considered the internal clock and scalar expectancy theory(3,34). In this model, saccade and repetition have been considered as distortion effects that can accelerate or decelerate the accumulation rate of pacemakers (clocks). This acceleration/deceleration can be multiplied to perceive time in absence of distortions (*t_Pure_*) by applying positive coefficients to adjust the rate of pacemakers (coefficients are higher than 1 for accurate time estimation and lower than 1 for inaccurate estimation). All the possible coefficients were applied to find the best performance of the model. A schematic/mathematic framework of this model has been presented in Figure 3.

The second model was implemented based on state dependent network models (6–8). To this end, a feedforward artificial neural network (ANN) with multi layer perceptron structure (two inputs perceptron, one hidden layer with 100 perceptron units in hidden layer and one output perceptron) was employed. A back-propagation algorithm was used for ANN training. We used 85% of dataset as training data and 15% of the dataset as test data. To evaluate the effect of different datasets on efficacy of ANN and to avoid overfitting, we used Bayesian regularization backpropagation algorithm that is a robust algorithm against overfitting problem. We consider the results of training and testing by looking at correlation value (R values) and RMSE between fitted values (by ANN model) and original values (from behavioral experiment). The mathematical processes of this modeling method are presented in Figure 3. We used artificial neural network toolbox in MATLAB 2019b to create the ANN model.

The third model was accomplished based on our previous hypothesis that state time in the brain has the same properties as physical time, represented by via vector units(30,31). In this model, time distortions are considered as different vector units to strengthen (dilatate) or destroy (compress) the vector of perceived time, or these effects can cancel in the absence of distortion effects (*t_Pure_*). This linear vector model was implemented by applying the cos(*ϕ, ψ*) (instead of the acceleration/deceleration weights of perceived time in scalar models). Where *ϕ, and ψ* were the assumed angles between the vectors of time distortion effects (i.e., saccade and repetition, respectively) and the veridical vector of time in the absence of time distortion effects. This model follows a linear formula where two distortion effects are linearly added to *t_Pure_* All the possible angles were tested, and the performance of model was obtained for these various angles. Figure 3 depicts this model mathematically and schematically. All the models have been generated by MATALB 2019b.

## 3 Results

### 3-1 Behavioral results

#### 3-1-1 Experiment 1

Ten participants performed this experiment, in counterbalanced blocks corresponding to three *tasks* (Figure 2-a). After analyzing saccade latency, one participant was excluded according to exclusion criteria. In each task, participants judged the variable duration (140, 170, 230, or 260ms) of a test stimulus (a horizontal or vertical grid) relative to the fixed-duration (200ms) of the last stimulus in a previously viewed series of 1 to 3 reference stimuli with the opposite (orthogonal) orientation (Figure 2-a). Participants were initially cued (by a small cross) to fixate their gaze just beyond one of four corners of the centrally located reference stimuli. In the main *Saccade Task*, this fixation cross shifted to the adjacent corner, triggering a vertical saccade with a stereotypical latency (268.4 ± 35.5ms). The test stimulus appeared just before saccade onset, such that saccades commenced at or near the end of the test stimulus, depending on its duration (Figure 2-b). Note that in this task, the test stimulus appeared at the same location as the reference stimuli but stimulated a different area of the retina (i.e., same spatial, different retinal location). To control for these spatial factors, we also included a *Fixation Task*, where the test stimulus appeared at the same location, but the fixation cross did not shift to induce a saccade (same spatial, same retinal location), and a *Retina-Matched Fixation Task*, where the test stimulus shifted like the fixation cross in the *Saccade Task* to stimulate the retinal location of the pre-saccadic stimuli (different spatial, same retinal). Thus, comparisons were performed among 6 conditions, i.e., Saccade/No-repetition (S-NoRep), Saccade/Repetition (S-Rep), Fixation/No-repetition (Fix1-NoRep), Fixation/Repetition (Fix1-Rep), and Retina-matched Fixation/No-repetition (Fix2-NoRep), Retina-matched Fixation/Repetition (Fix2-Rep). Furthermore, interaction of repetition and task was evaluated.

To help distinguish the effects of saccades (time underestimation) and repetition (overestimation) we divided the data into incorrect response for shorter versus longer trials. Figure 4-a shows the percentage of incorrect responses for shorter trials (overestimation) in different conditions (individual responses were presented by color dots and distributions of responses were presented by violin plots). FDR revealed significant differences between repetition and no-repetition conditions in all tasks. This suggests repetition causes overestimation of time either in fixation or saccade tasks. No significant interaction was observed between repetition effect and tasks. Statistical details are presented in the caption of figure 4.

**Figure 4:**
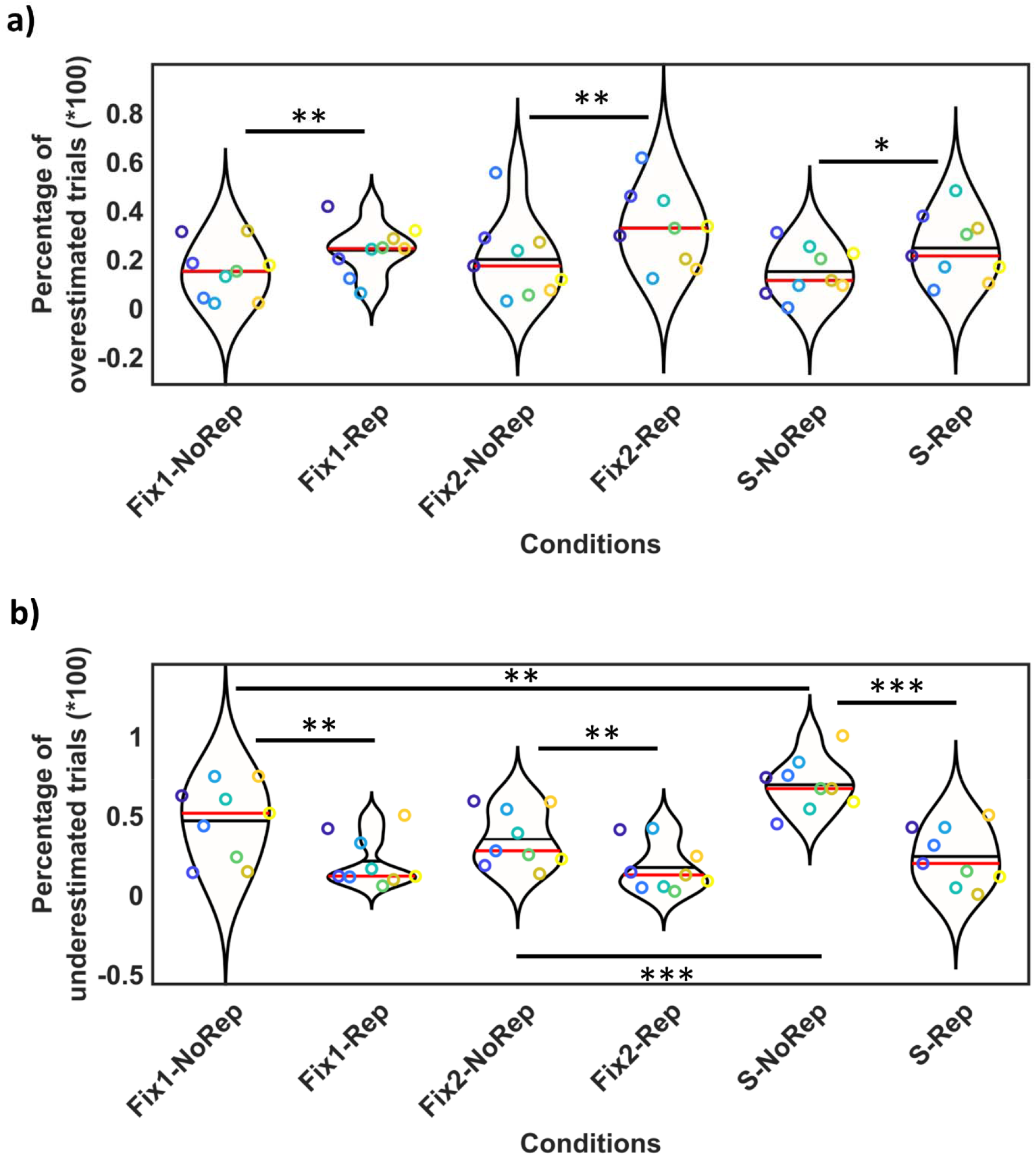
Statistical results of the first experiment, showing significant results between different tasks and presentations of test stimulus. The distributions of responses were presented by ‘violin plot’ fits (where the horizontal axis shows different conditions, and the vertical axis shows % overestimated or underestimated trials). Individual responses were presented by color dots (responses of each participant can be tracked by a specific color and a specific vertical location in each condition). One-star (*) indicates (p<0.05), two-stars (**) indicates (p<0.01), and three-stars (***) shows (p<0.001). Data are divided into incorrect responses for shorter versus longer trials: a) Percentage of incorrect responses for shorter trials (overestimation of time). Significant differences were observed between: Fix1-NoRep vs. Fix1Rep (t=-3.55, p=0.0075), Fix2-NoRep vs. Fix2Rep (t=-3.79, p=0.0053), S-NoRep vs. S-Rep (t=-3.14, p=0.013). b) Percentage of incorrect responses for longer trials (underestimation of time). Significant differences were observed between 1) repetition vs. no-repetition: Fix1-NoRep vs. Fix1-Rep (t=4.95, p=0.001), Fix2-NoRep vs. Fix2-Rep (t=4.62, p=0.002), S-NoRep vs. S-Rep (t=11.23, p<0.001), and tasks: S-NoRep vs. Fix1-NoRep (t=-3.63, p=0.007), S-NoRep vs. F2-NoRep (t=-7.49, p<0.001).

Incorrect responses for longer trials (underestimation) are presented in figure 4-b. In this figure individual responses were presented by color dots and distribution of responses were presented by violin plots. FDR revealed significant decreased underestimation in repetition conditions in comparison to repetition conditions in all tasks. Comparisons between tasks revealed significant time underestimation in the saccade condition (in comparison to both fixation conditions). No significant differences were observed between two eye-fixed conditions. Furthermore, we observed significant interactions between repetition effects and tasks (*S-Rep* minus *S-NoRep*) vs. (*Fix1-Rep* minus *Fix1-NoRep*) (t=-3.12, p=0.014) and (*S-Rep* minus *S-NoRep*) vs. (*Fix2-Rep* minus *Fix2-NoRep*) (t=-5.06, p<0.001)). Post hoc analysis revealed that there is no significant difference between S-Rep, Fix1-Rep, Fix2-Rep conditions. It suggests that repetition cancels the underestimation effect of saccades. Statistical details are presented in the caption of figure 4.

These results show that overestimation of time was always present after repetition of the reference stimulus, and that this effect superseded the time compression effect in the presence of a saccade. The underestimation of time for individual participants was always observed in the saccade-norepetition condition whereas when the reference stimulus was repeated no significant underestimation was observed in the saccade task.

#### 3-1-2 Experiment 2

21 participants performed this experiment. Three participants were excluded because of exclusion criteria (inappropriate saccade latency). This experiment was similar to the first experiment, except two tasks (fixation and saccade) (instead of three tasks) were performed by participants and timing was different (duration of test was 70ms and reference stimulus duration was one of the variable durations: 30, 50, 90, 110ms). Furthermore, participants performed horizontal saccades instead of vertical. In the *Saccade Task*, latency of saccade was 263.7 ± 41.4ms.

Figure 5-a presents the percentage of incorrect responses for shorter trials (overestimation of time) in different conditions (individual responses were presented by color dots and distributions of responses were presented by violin plots). The FDR analysis revealed that repetition significantly increased incorrect responses in both tasks that means pre-stimulus repetition cause to overestimation of time either in fixation or saccade tasks. No significant difference was observed between fixation and saccade tasks. Furthermore, interaction between repetition effect and tasks was not significant. Statistical details are presented in the caption of figure 5.

**Figure 5:**
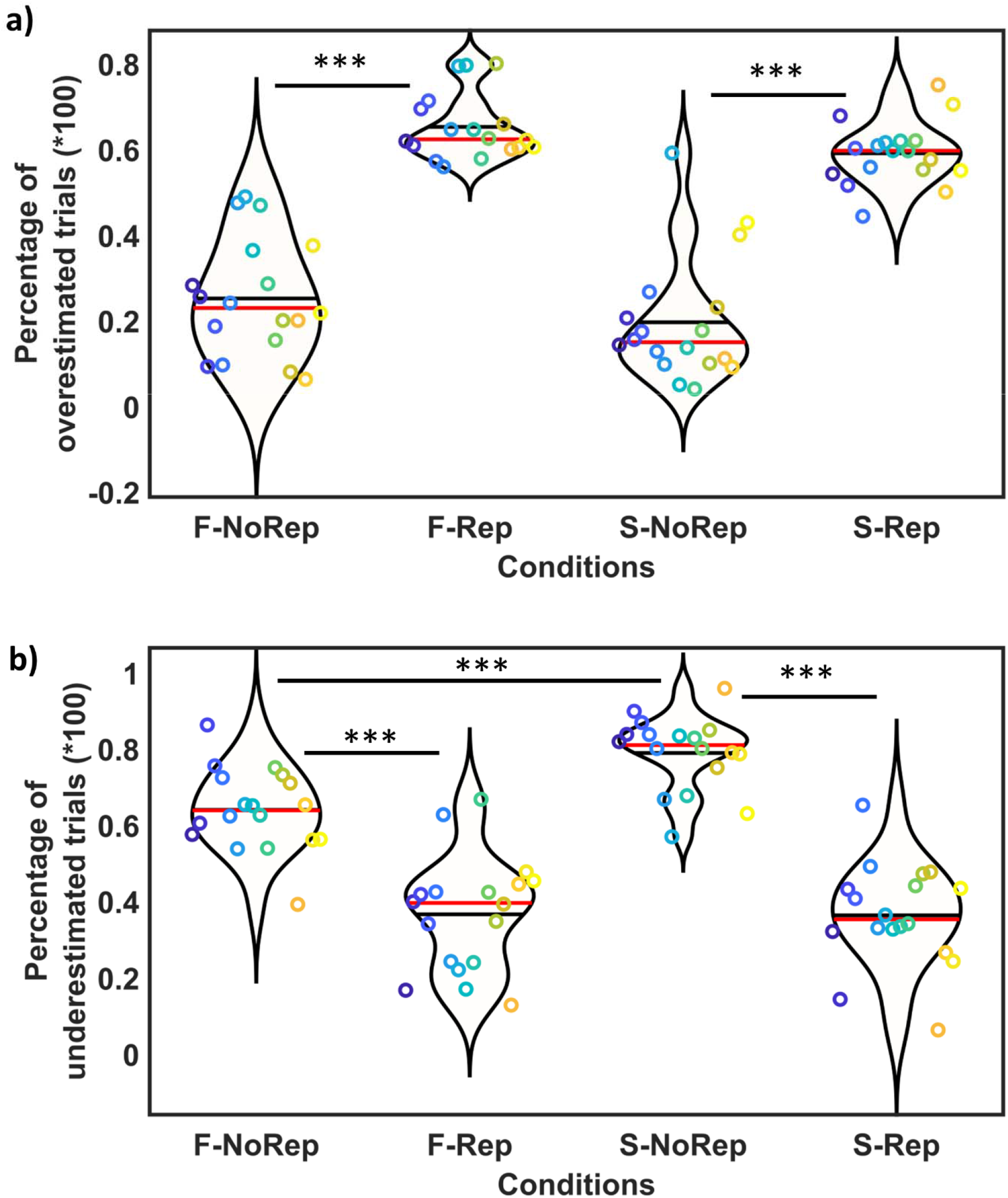
Statistical results of the first experiment, showing significant results between different tasks and presentations of test stimulus. The distributions of responses were presented by ‘violin plot’ fits (where the horizontal axis shows different conditions, and the vertical axis shows % overestimated or underestimated trials). Individual responses were presented by color dots (responses of each participant can be tracked by a specific color and a specific vertical location in each condition). One-star (*) indicates (p<0.05), two-stars (**) indicates (p<0.01), and three-stars (***) shows (p<0.001). Data are divided into incorrect responses for shorter versus longer trials: a) Percentage of incorrect responses for shorter trials (overestimation of time). Significant differences were observed for repetition effect: F-NoRep vs. F-Rep (t=-12.40, p<0.001), S-NoRep vs. S-Rep (t(17)=10.71, p<0.001). b) Percentage of incorrect responses for longer trials (underestimation of time). Significant differences were observed between 1) repetition vs. no-repetition: F-NoRep vs. F-Rep (t=6.93, p<0.001), S-NoRep vs. S-Rep (t=10.02, p<0.001), and tasks: S-NoRep vs. F-NoRep (t=-4.57, p<0.001)]

Figure 5-b shows incorrect responses for longer trials (underestimation of time). In this figure, individual responses were identified by color dots and distribution of responses were presented by violin plots. FDR revealed significant time underestimation in the saccade/no-repetition condition in comparison to fixation/no-repetition condition. Significant differences were also observed between repetition and no-repetition conditions in both tasks (i.e., S-NoRep vs. S-Reo and F-NoRep vs. F-Rep). Significant interaction was observed for repetition effect and task (t= 2.56, p=0.021). Post hoc analysis showed that there is no significant difference between SR and FR conditions. This later result suggests that repetition cancels the underestimation of time in saccade condition.

Consistently, this experiment showed that repetition of a stimulus before the test stimulus led to overestimation of perceived duration for test stimulus. Also, saccade compressed the perceived duration of test stimulus that was presented 100ms before saccadic eye movement. Like the first experiment, results of the second experiment confirm that this saccadic time compression is canceled when a repeated series of stimuli is presented before the saccade.

### 3-2 Modeling results

In the second part of this study, we defined individual timing performance (ITP) by comparing the individual time judgments in trials with time distortion effects (i.e., repetitions and saccade) and trials without distortion effects. We integrated behavioral results from both experiments and used the calculated ITPs (for both experiments and longer or shorter trials) to test between the models shown in Figure 1. To test the ability of each model to explain our observed Saccade-Repetition (SR) interactions, the Saccade-NoRepetition condition (S), and Fixation-Repetition condition (FR) were used as inputs to the model and the outputs were responses in the Saccade-Repetition condition (SR), i.e., each model was required to integrate the separate saccade and repetition effects in order to reproduce the combined effect. Each model was trained or optimized to best fit the actual output data. Figure 6 illustrates the goodness of fit of each model against the data as scatter plots (model output as a function of actual Saccade-Repetition data), and beneath these, the corresponding residuals of fit (i.e., Euclidean distances between outputs of models and the data). The inputs are derived from *ITP_R_* and *ITP_S_ in* two experiments and output for this set was *ITP_SR_*. In the first experiment, since we had two fixation conditions, *ITP_S_* and *ITP_R_* were calculated for *fixation/saccade*, and *retina-fixed/saccade* condition combinations. This ITP set were separately calculated for longer and shorter trials. Then, in this experiment, four *ITP* set (*fixation/saccade-*longer trials, *fixation/saccade-shorter* trials, *retina-fixed/saccade-longer* trials, and *retina-fixed/saccade-shorter* trials) were obtained for each participant (for all nine participants we had totally 4×9=36 ITPs). For the second experiment, ITP set of 18 participants were calculated for *fixation/saccade* trial combination. For each participant two ITP set were obtained according to their responses for *longer* and *shorter* trials. Then, total number of ITPs were 2×18=36 in this experiment.

**Figure 6:**
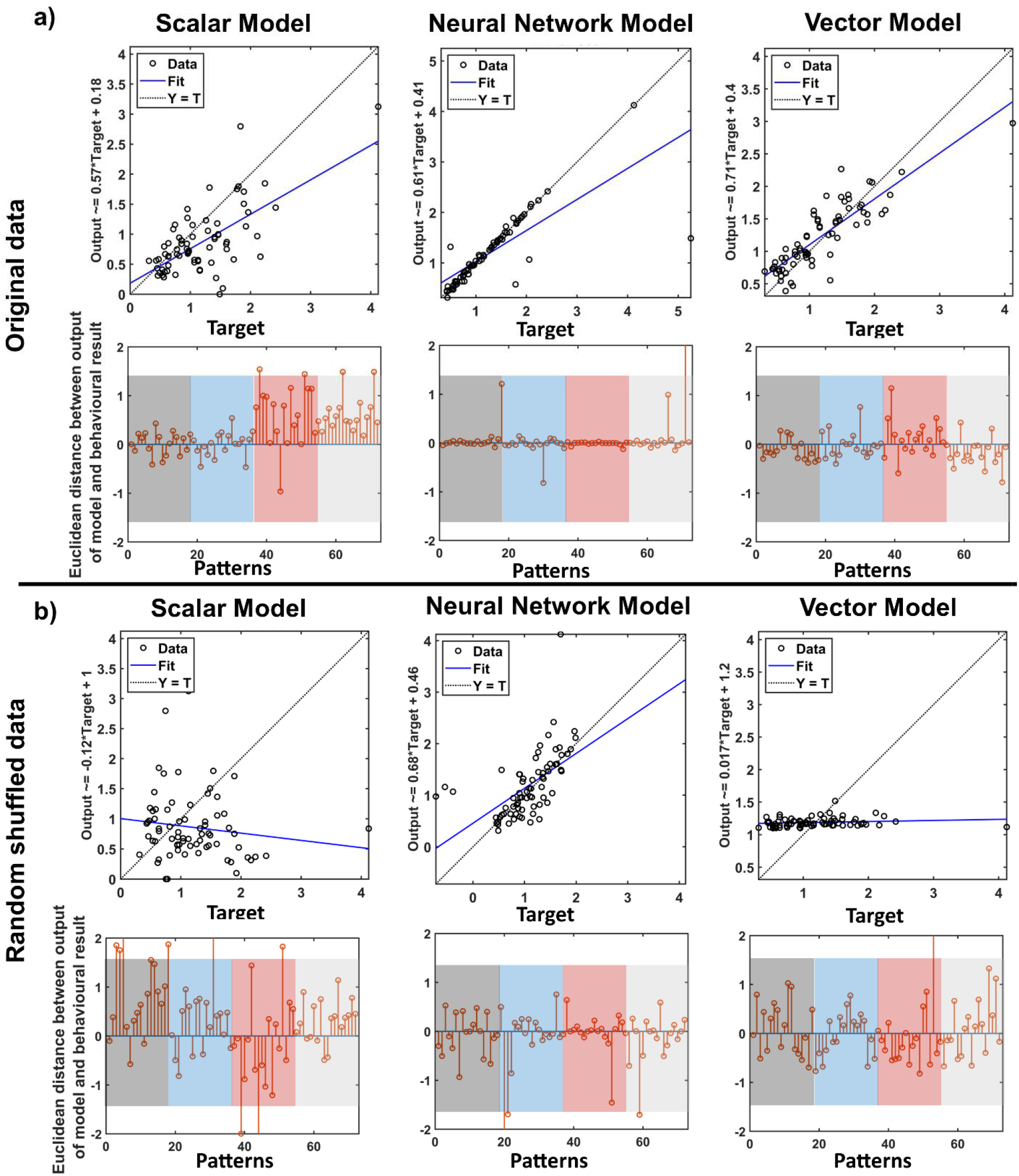
The correlations and residuals (Euclidean differences between output of models and behavioural results) showing goodness of fit for each model, where high correlation and low residuals signify good fit and higher residuals and lower correlation signify relatively poor fits. **(a):** correlation and residuals for fits to actual data. The residuals are sorted according to the experiments and trials: longer trials in second experiment (1 to 18; gray shadow), shorter trials in second experiment (residuals19 to 36; blue shadow), longer trials in first experiment (residuals 37 to 54; red shadow), shorter trials in first experiment (residuals 55 to 72; grey shadow). (b): residuals between output of models and random shuffled patterns (across participants and states) of behavioural results (outputs were assigned to irrelevant inputs (see Methods for details). The colors are similar as the part (a). The *Scalar Model* (left column) yields moderate fits for original data, the *Neural Network Model* (centre column) yields good fits, but for both the original and shuffled data, whereas the *Vector Model* (right column) yields the best fits (i.e., lowest residuals and high correlation) and only for the original data.

According to the third criterion, a good model should be valid against different task conditions (e.g., different durations), then we integrated all ITPs from two experiments and both longer and shorter trials. Totally 72 ITP sets were used as input/output samples for models. Figure 6-a shows fits made to the original data whereas the part (b) shows fits made to shuffled data (see methods) as a control. To quantify the goodness of fit for each model, we performed regression analysis (Table 2) that accounted for the different combinations of input used for testing and participant number as factors and took the Root Mean Square Error (RMSE) of the residuals. The best model should show good fits that are specific, i.e., high correlations and low residuals, and only to the original data, not the shuffle control for all samples (regardless of experiment or type of trials).

**Table 2:** the performance of different models to predict the behavioural results in four different i.e., condition 1) one repetition in *Fixation Task × Saccade Task*, condition 2) two repetitions in *Fixation Task × Saccade Task*, condition 3) one repetition in *Retina-Matched Fixation Task × Saccade Task*, and condition 4) two repetitions in *Retina-Matched Fixation Task × Saccade Task*. The R^2^ and p-value were obtained from the multiple linear regression analysis. RMSE was calculated based on the Euclidean distance between output of the models and behavioral data.

| **Dataset** | **Scalar model** | **Vector model** | **ANN model** |
| --- | --- | --- | --- |
| **Original data** | R=0.64, p-value<0.01<br>RMSE=0.56 | R=0.86, p-value<0.01<br>RMSE=0.32 | R=0.78, p-value<0.01<br>RMSE: 0.49 |
| **Random shuffled data** | R=-0.13, p-value=0.26<br>RMSE: 0.93 | R= 0.15, P-value=0.22<br>RMSE=0.61 | R=0.57, p-value<0.01<br>RMSE: 0.54 |

After Bonferroni correction, all three models showed significant correlations between fitted outputs and original data. high correlation values (R-values>0.7) were obtained using the vector and ANN models. The scalar model showed moderate correlation (R-value=0.64). The highest accuracy (largest R-value and smallest RMSE) was achieved using linear vector model (table 2). This showed that the vector and ANN models are reliable models to describe both distortion effects in different condition. But the scalar model is in moderate range of sensitivity. Figure 6-a showed that scalar model was particularly reliable only in one experiment.

As shown in figure 6-b and table 2, when we tested three models against random-shuffled data to examine specificity of models, two models (the scalar and vector models) showed nonsignificant correlations between fitted outputs and random-shuffled outputs. However, ANN exhibited significant moderate correlation (R=0.57). This suggest that the scalar and vector models are specific models for time perception, but ANN is not a specific model.

In summary, the performance of scalar model was in moderate range, the neural network model performed better but was not specific to the actual dataset, whereas the linear vector model fit both of our criteria: it performed well and was specific to the original unshuffled datasets for both experiments.

## 4 Discussion

In this study, we asked how two separate time distortion effects (repetition time dilation and perisaccadic time compression) interact, and which time perception model can best explain the derived results. Overall, the behavioral results confirm previous findings on perisaccadic time compression(14,35) and relative effect of time dilation after a series of repetitive stimuli(20,21,24) but expand on this by showing how they interact. Results showed the relative time dilation induced by repetition of a reference stimulus before test stimulus neutralized the presence of an intervening saccade (i.e., the interactive effect of perisaccadic time compression and repetition time dilation is similar to single effect of repetition time dilation alone). To model this interaction, the resulting data were used as inputs/outputs for models of time perception (scalar clock, non-linear artificial neural network, linear vector). The linear vector model performed best, both in terms of specificity and ability to fit the data. Here, we will consider the experimental results in more detail, and discuss their implications for broader time perception theories.

In agreement with previous investigations(12,14,15,35), our results showed that saccades compress the subjective perception of the duration of a briefly presented visual stimulus. Whereas previous investigations used empty temporal intervals (i.e. the interval between presentation of two stimuli(14) has been judged instead of duration of one presented stimulus), we presented the test stimulus during perisaccadic interval and participants judged the duration of the stimuli themselves. This demonstrates that, notwithstanding the presence of different mechanisms for time perception of brief empty or filled intervals(36), saccadic time compression occurs in both cases. On the other hand, our results support previous studies(13,20,21,24) that found the relative duration of a novel stimulus after a series of repeated stimuli is perceived to be longer than the last repeated stimulus (in the repetitive series) with the same objective duration.

In *the Retina-Matched Fixation Task* (first experiment), the test stimulus was presented in a different spatial location (but same retinal location) relative to the repeated reference stimulus. Thus, in this task, the test stimulus shows two types of novelty compared to the reference stimuli: different location and different orientation. However, in this condition time dilation was not significantly different from that in our *Fixation Task* (where the test stimulus was presented in the same objective and retinal location as the reference stimulus). This suggests that objective spatial location did not have a strong effect on perceived time. This might be because the new location of our test stimulus was always predictable, and one could predict that the test stimulus would be presented in another possible location that was different from the location of reference stimulus. Whereas the randomized number of repetitions was not predictable in all tasks.

Various, often disconnected explanations have been provided for these phenomena. Saccadic remapping(14,37), transient responses in visual cortex(23), saccadic suppression(16,27,37,38) and attention(16) have been assumed to be associated with perisaccadic time compression, whereas time dilation of a novel stimulus (after a series of repeated stimuli) has been explained by attention(24) and repetition suppression mechanisms(13,27). The latter two explanations (neural suppression and attention) are common to both phenomena (saccadic time compression and time dilation after a series of repeated stimuli) but the other proposed mechanisms for saccadic suppression can not explain time dilation for the novel stimulus following a repeated stimulus. Therefore, when considering how these phenomena might interact, we will focus on attention and neural suppression.

We will first consider the joint effects of attention on time distortions induced by saccade and repetition(16,24), because this can be directly related to the internal time clock model simulated above. In the internal clock model, attention is considered as a switch between the pacemaker and accumulator(3,5,39) (Figure 1-a). Higher levels of attention allocation can turn on this switch, thus causing the accumulator to collect more timing pulses and overestimate time(24,39). Alternatively, a recent saccade / time perception study suggested the decreased attentional allocation to the stimulus leads to decreased time pulse accumulation in the internal clock, thus resulting in perisaccadic time compression(16).

Based on this model, subjective time is a scalar parameter(3) that has accumulative property. This model thus predicts that a scalar combination of saccade and repetition trials should correlate with saccade-repetition trials (Figure 1-a and figure 3). However, this did not clearly occur in our results: instead, repetition obliterated the saccades effect. Thus, when these data were used to simulate the internal clock model, moderate correlations resulted.

The idea that suppression of brain activity can compress time perception was originally presented by Eagleman and Pariyadath as an *energy model* in time perception(12,19–21,27). The basic idea is that just before and during saccade execution (the perisaccadic interval) certain brain areas show reduced activity(37,40–42), and thus energy expenditure(12,13,27). Since stimulus novelty (especially after a repeated stimulus) can increase neural activity / energy expenditure, it can (according to this theory) also dilate perceived time(13,27) (see supplementary figure 1). A limitation of this model is that the causal link from energy metabolism *to* information processing is indirect, at best. However, taking this theory at face value, when both effects (repetition and saccade) occur simultaneously, the level of energy spent in the brain will be decreased by one factor (the saccade) and increased by other (novelty after repetition). Again, this can be represented mathematically as a combination of two energies (two coefficients) that are spent in different brain regions. Based on this assumption, behaviorally, we expect that one effect would cancel or weaken another effect. The correlation between outputs of the scalar model and empirical data was significant but R-value remained in the moderate range. Thus, in this case, scalar models of time perception (e.g., internal clock, and energy models) were partially able to describe the joint time distortion effect of repetition and saccades.

Since the scalar timing models showed results with moderate accuracy to justify the joint time distortion effects of saccade and repetition, a SDNs model was employed as an alternative method. This model explains state-dependent computations in neural network level presents subjective time(7,8,43–45). Previous SDNs modeling studies have showed that artificial neural networks can simulate the behavioural responses in different timing conditions such as rhythm perception(8) and can encode time-varying sensory and motor patterns(46). This approach is similar to other artificial neural network studies that try to model a connectivity structure between incoming visual/auditory stimuli and behavioural responses. By this way, non-temporal perceptual classification approaches (e.g. feedforward neural networks) that are usually working based on Hebbian law, can be used to classify and predict subjective time(29). Consistent with the above studies, our, artificial neural network was able to fit the data in our study. However, such a network can fit almost anything, including a random shuffle of our data, i.e., it may have succeeded through overfitting. Thus, while this model was able to retroactively predict our data, the prediction was not specific.

A common element between scalar timing and SDN models is that they do not explicitly consider the concept of time in modern physics (time in Minkowski space). Objective time in these models is supposed to be the sequence of events, leading to subjective time perception. There is an interesting similarity between internal clock model and Newtonian absolute time, because both frameworks assumed an absolute generator that can create passage of time (Table 1). On the other hand, the SDN models assume moments can be coded via a nontemporal classification network and a Hebbian law is involved in learning and recognition of temporal patterns. However, based on modern physics theories each observer measures time as an intrinsic property from the environment, and this property can be changed based on information (entropy) and speed of both the environment and observer(32,47). If we consider the brain as a physical observer, the same timing parameters should be hold in the brain and in the physical universe (Table 1). As time is characterized as a vector in Minkowski space, we used this concept in our last modeling approach. This vector model was originally suggested for time perception in long durations(30) but it can be proposed as a general time perception model for all intervals(31).

These time vectors can be defined by entropy and speeds in a physical system. In physiological terms, entropy can be conceptualized through different hypothesis. For example, order/disorder of states in dynamical functional connectivity patterns can change information entropy in functional brain networks and this alteration may compress or dilate perceived time(30). From a thermodynamical perspective, fluctuation of brain tissue temperature (as a thermodynamical system that exchanges heat with its surrounding) changes brain entropy and likewise may alter perceived time(31).

Our modeling results show that the vector model(30,31) exhibited both sensitive and specific simulations of joint saccadic and repetition time distortion effects, i.e. in this model separate saccade and repetition result inputs predicted the combination of both effects. Furthermore, this model presented accurate results for both experiment (regardless of duration of intervals and type of trials). By representing perceptual effects as vectors and predicting the angles between these (Figure 3), this model was able to predict their interactions accurately and specifically. Conversely, in this model different time distortion effects can be considered as vectors in different directions that are added/subtracted to/from the original time vector (A detailed mathematical description of the specific vector interactions in our model can be found in *Supplementary Discussion*).

Overall, the results of vector timing model confirmed our hypothesis that subjective time has deep similarities to physical time and satisfied the first postulate of the special relativity theory(47). In this model, more information than magnitude (that is considered in scalar models) is thought to be required to model dilation and compression of subjective time. Then, subjective time is characterized by vector units of time with two properties: magnitude and direction. When two time distortion effects are simultaneously interfered subjective time, two vectors with different magnitudes and directions are added to subjective time. Presumably, this information theory model is instantiated at the level of neural networks and cellular signals, but at this time the biological mechanism is unknown.

In conclusion, we show here that global theories of time perception can be used to predict interactions between seemingly disparate experimental phenomena and conversely, those such interactions can help test between global theories. In particular, this study is the first to show the interaction between saccadic time suppression and repetition time dilation, and that this interaction follows the specific predictions of the vector model of time perception. In terms of broader implication, this is the first empirical-theoretical investigation that directly shows subjective time can be represented by the same time parameter as that used in physics.

## Acknowledgements

We thank Drs. P. Cavanagh, V. Bharmauria and P.A. Khoozani for editorial comments on the manuscript, and Dr X. Yan, S. Sun, and G. Tomou for technical support. This project was funded by a Natural Sciences and Engineering Research Council of Canada (NSERC) Discovery grant. A.H. Ghaderi was supported by the Vision: Science to Applications Program, funded in part by the Canada First Research Excellence Fund. J.D. Crawford was supported by a Canada Research Chair.

