## Supplementary materials for "Linear vector models of time perception account for saccade and stimulus novelty interactions"

Table of contents


| TITLE | PAGE |
|---|---|
| SUPPLEMENTARY DISCUSSION | 1 |
| SUPPLEMENTARY FIGURE 1 | 2 |
| REFERENCES | 2 |

### 1) Supplementary discussion:

According to the results of this study, the vector characterization of perceived time can explain various aspects of perisaccadic time distortion effect. For example, the results show that the predicted angles between perisaccadic time distortion effect ($t_S$) and pure time ($t_{pure}$) can be higher than 90 degree (see supplementary figure 3-a). Mathematically, when the magnitude of perisaccadic time distortion ($t_S$) is bigger than $t_{pure}$ and both vectors lay in opposite directions, then the output of this vector operation is less than zero (see Supplementary Figure 4 to find the mathematical details). As the result, time will be perceived inversely (backwards or toward the past)! This strange condition (inverse time) has been reported in previous studies[1,2] during perisaccadic interval when two successive events are presented in very short intervals (less than 70 ms) and participants were asked to judge the order of events[2,3]. Most of the participants reported an inverse order for the presented events[2,3]. This effect has not been explained by previous global models in time perception (e.g. scalar timing models). Obviously, if one consider perceived time as the collected pulses by an accumulator (a scalar perspective about perceived time), this effect can not be explained (because we can not assume negative pulses of time distortion effects to negate the feedforward time). But the introduced vector model of time perception simply predicts this effect. Furthermore, the best fit between the outputs of our vector model and the empirical data determined the arbitrary parameters of the vector model (i.e. angles) in somehow to predict time dilation for the repetition effect ($t_R$).

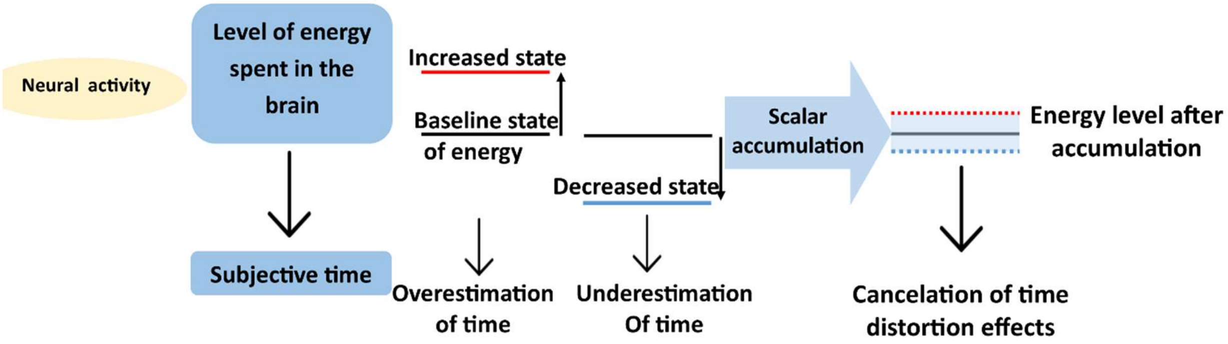


**Supplementary Figure 1:** *Energy model:* In this model, the amount of energy spent in the brain (directly related to neural activity) is assumed to be involved in time perception. Higher values of this scalar parameter (energy spent) cause to overestimation of time and voice versa. When energy of brain affected by two opposite effects (one decreases the neural activity and another increase that) the combination of energies will be near to baseline of energy (energy of brain before the distortion effects) and time will be perceived without distortion.


1. Binda, P., Cicchini, G. M., Burr, D. C. & Morrone, M. C. Spatiotemporal Distortions of Visual Perception at the Time of Saccades. *J. Neurosci.* **29**, 13147–13157 (2009).
2. Morrone, M. C., Ross, J. & Burr, D. Saccadic eye movements cause compression of time as well as space. *Nat. Neurosci.* **8**, 950–954 (2005).
3. Terao, M., Watanabe, J., Yagi, A. & Nishida, S. Reduction of stimulus visibility compresses apparent time intervals. *Nat. Neurosci.* **11**, 541–542 (2008).
